# Establishing Cerebral Organoids as Models of Human-Specific Brain Evolution

**DOI:** 10.1101/500934

**Authors:** Alex A. Pollen, Aparna Bhaduri, Madeline G. Andrews, Tomasz J. Nowakowski, Olivia S. Meyerson, Mohammed A. Mostajo-Radji, Elizabeth Di Lullo, Beatriz Alvarado, Melanie Bedolli, Max L. Dougherty, Ian T. Fiddes, Zev N. Kronenberg, Joe Shuga, Anne A. Leyrat, Jay A. West, Marina Bershteyn, Craig B. Lowe, Bryan Pavlovic, Sofie R. Salama, David Haussler, Evan E. Eichler, Arnold R. Kriegstein

## Abstract

Direct comparisons of human and non-human primate brain tissue have the potential to reveal molecular pathways underlying remarkable specializations of the human brain. However, chimpanzee tissue is largely inaccessible during neocortical neurogenesis when differences in brain size first appear. To identify human-specific features of cortical development, we leveraged recent innovations that permit generating pluripotent stem cell-derived cerebral organoids from chimpanzee. First, we systematically evaluated the fidelity of organoid models to primary human and macaque cortex, finding organoid models preserve gene regulatory networks related to cell types and developmental processes but exhibit increased metabolic stress. Second, we identified 261 genes differentially expressed in human compared to chimpanzee organoids and macaque cortex. Many of these genes overlap with human-specific segmental duplications and a subset suggest increased PI3K/AKT/mTOR activation in human outer radial glia. Together, our findings establish a platform for systematic analysis of molecular changes contributing to human brain development and evolution.

## Introduction

The three-fold expansion of the human cerebral cortex is one of the most conspicuous features distinguishing humans from other great apes (Herculano-Houzel, 2012). This size difference is already apparent during early cortical development at mid-gestation, prior to the completion of neurogenesis (Sakai et al., 2012). Longstanding models propose that increased numbers of neural stem and progenitor cells could account for human brain expansion (Rakic, 1995). Radial glia act as neural stem cells to generate excitatory neurons of the cerebral cortex (Miyata et al., 2001; Noctor et al., 2001), and recent comparative studies suggest a cell-intrinsic increase in proliferative divisions among radial glia as a candidate developmental mechanism for human brain expansion (Otani et al., 2016). Nonetheless, the molecular basis for differences in developmental cell behavior remains poorly understood because primary brain tissue is largely inaccessible from chimpanzees, our closest living relatives, during developmental stages in which neurons of the cortex are generated.

Induced pluripotent stem cells (iPSCs) derived from great apes provide a platform for experimentally addressing how human-specific genetic changes differentially affect aspects of development (Marchetto et al., 2013; Enard, 2016; Gallego Romero et al., 2015; Prescott et al., 2015; Silver, 2016). Organoid models derived from pluripotent stem cells harness natural properties of self-assembly to mimic early developmental processes across diverse tissues (Eiraku et al., 2008; Eiraku et al., 2011; Clevers, 2016). Recent studies have begun to compare gene expression between human and chimpanzee cerebral organoids from a limited number of individuals (Mora-Bermúdez et al., 2016), but systematic quantitative comparisons across multiple individuals and time-points are required to distinguish species differences from individual differences. Additionally, primate outgroups beyond chimpanzee are necessary to determine which evolutionary changes are derived in the human lineage. Similarly, analysis of the fidelity of organoid models to primary tissue requires comparing key sources of biological and technical variation across a range of protocols and individuals.

Single cell RNA sequencing provides an opportunity to compare gene expression in homologous cell types between primary tissue and organoid models and across species (Pollen et al., 2014; Camp et al., 2015). Although developing tissue and organoid models contain a diversity of cell types, new analysis methods allow for the identification of homologous cell types and gene regulatory networks based on the expression of thousands of genes in single cells (Butler et al., 2018; Nowakowski et al., 2017). Here, we use single cell gene expression comparisons across the span of cortical neurogenesis to undertake three analyses that together enable the study of gene regulatory evolution during human brain development (Figure 1A). First, we examine the extent to which cell types, gene co-expression patterns, and developmental trajectories from primary tissue samples are preserved in organoid models. Second, we explore how gene expression patterns diverge between human and macaque during cortical development using primary tissue. Finally, we analyze which of the expression differences between human and macaque emerged along the human lineage in the last six million years using human and chimpanzee organoid models that give us an otherwise inaccessible window into patterns of early brain development in our closest living ancestor.

**Figure 1.**
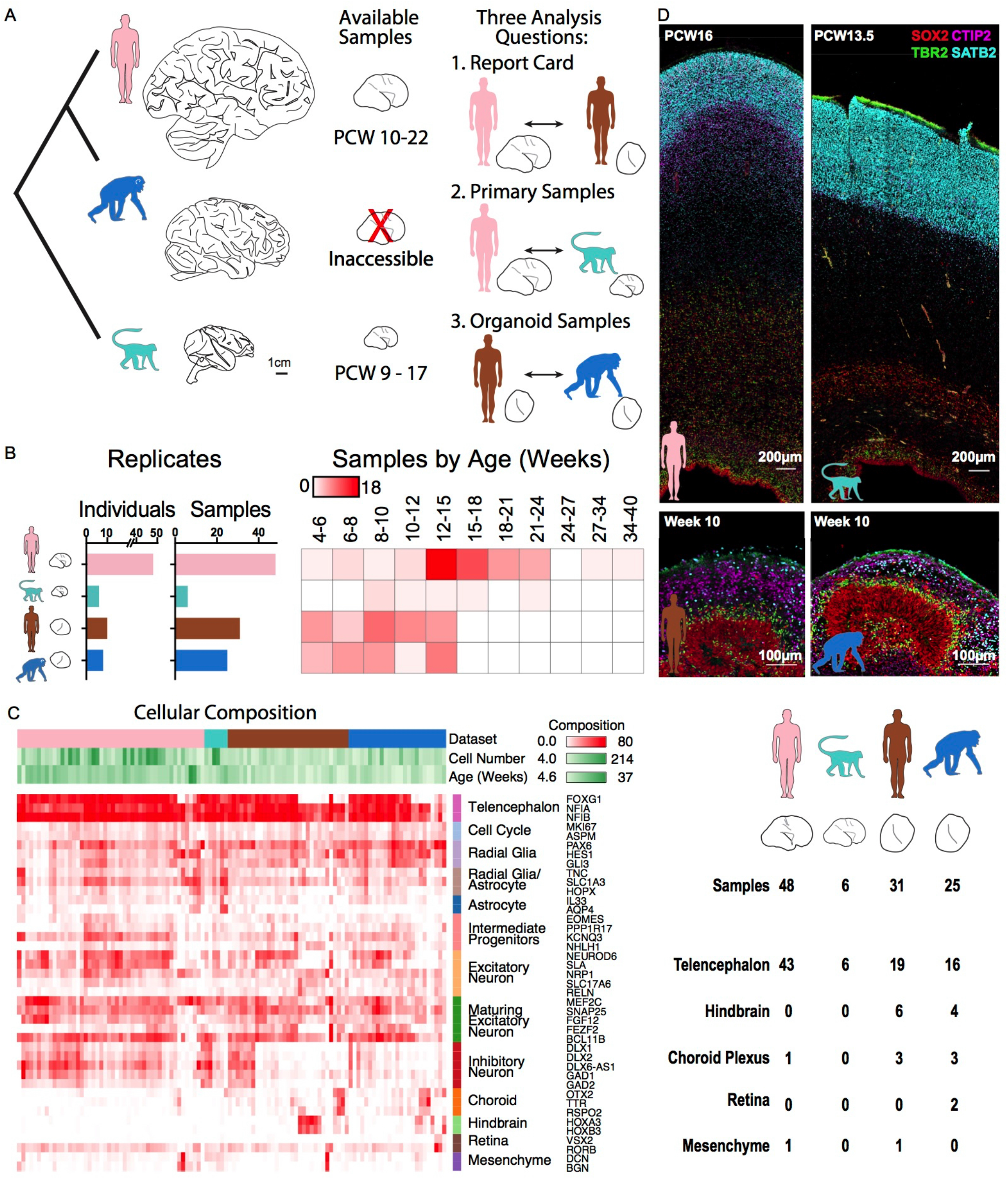
Organoid models reflect normal features of human and chimpanzee brain development. **a**) The human brain has expanded dramatically compared with other primates, but brain tissue is largely inaccessible from developing chimpanzee. To compare human and chimpanzee development, our study focuses on three analysis questions that integrate data from primary human (pink) and macaque samples (light blue) with human (brown) and chimpanzee (blue) organoid models. **b**) Histograms and heatmap depict the number of individuals and primary or organoid samples and the distribution of samples over post conception or post differentiation weeks. **c**) Heatmap represents the fraction of cells expressing each marker gene across cells from each primary sample or organoid, and table summarizes the number of samples that predominantly express markers for a given regional identity. **d**) Immunohistochemistry for markers of radial glia (SOX2), intermediate progenitors TBR2 (EOMES), and neurons CTIP2 (BCL11B), SATB2, reveals histological and cellular features of normal neurogenesis in the germinal zones of human and chimpanzee cerebral organoids, but a much more extensive intermediate zone and cortical plate in primary samples.

We find that organoid models preserve the vast majority of gene co-expression patterns observed in primary tissue during cortical development, supporting the utility of these models for studying the evolution of gene regulation. Nonetheless, across commonly used protocols, organoid models display upregulation of glycolysis, endoplasmic reticulum stress, and electron transport pathways, isolating potentially important differences with primary tissue, while also highlighting avenues for further improving this useful model. Similarly, we find striking conservation of gene networks across human, chimpanzee and macaque. However, we also identify a common set of 261 genes that are differentially expressed between human and macaque primary cortical cells and between human and chimpanzee organoid cells. These candidate human-specific gene expression differences are largely distinct from those reported in fibroblasts and pluripotent stem cells (Gallego Romero et al., 2015) and include many genes overlapping recent segmental duplications. Interestingly, we find increased activation of the PI3K/AKT/mTOR pathway specifically in human outer subventricular zone radial glia, suggesting a molecular mechanism that may contribute to differences in human neural stem cell behavior.

## Results

### Cell diversity in primary samples and organoid models

Cortical development involves diverse communities of cells, including excitatory neuron lineage cells (radial glia, intermediate progenitor cells, excitatory neurons), inhibitory neurons that migrate to the cortex from the ventral telencephalon, glial cells (astrocytes, oligodendrocytes, and microglia), and vascular cells. Organoid models may capture a subset of these cell types while also introducing additional sources of variation in cell types, cell states, and gene expression (Quadrato et al., 2017). We reasoned that single cell gene expression would allow us to evaluate the extent to which cell types, gene regulatory networks, and developmental trajectories are preserved in organoid models. Therefore, we designed a large-scale experiment to compare single cell gene expression using a standardized approach across primary tissue from 48 human samples (Nowakowski et al., 2017) and 6 macaque samples with 56 organoids derived from 10 human and 8 chimpanzee individuals, distributed across stages of cortical neurogenesis (Figure 1B). We generated organoids based on a protocol developed in the Sasai lab (Kadoshima et al., 2013), and we further analyzed published single cell gene expression data from two other organoid protocols representing a continuum of guided to unguided patterning (Camp et al., 2015; Kadoshima et al., 2013; Lancaster et al., 2013; Sloan et al., 2017) (Figure S1). To analyze the largest number of homologous genes across human, chimpanzee, and macaque, we updated gene models and orthology assignments in both chimpanzee and macaque genomes using the comparative annotation tool kit (Fiddes et al., 2018), providing a more accurate and comparable estimate of gene expression levels between species. In addition, we recently improved the contiguity of the chimpanzee genome through de novo assembly, increasing alignment rates (Kronenberg et al., 2018). Using these improvements, we were able to align reads to each species’ native genome and examine gene expression across 49,360 orthologous genes (Figure S1). The number of genes detected per cell was comparable in human and chimpanzee organoids, but slightly lower in macaque, indicating that expression values may be underestimated for some macaque genes (Figure S1).

To characterize the cellular heterogeneity of organoid models, we first analyzed the frequency of cells expressing marker genes for cortical cell types as well as for common off-target lineages. The majority of human and chimpanzee organoids expressed markers of telencephalic regional identity, including cortical excitatory lineage cells, and a subset of inhibitory neurons from ventral telencephalon (Figures 1C, S1). A minority of organoids predominantly contained off-target lineages, including hindbrain, choroid plexus, retina, and mesenchymal cells, with a few individuals mainly accounting for the bias in differentiation potential (Figure S1). We observed a similar proportion of cortical lineage cells and off-target lineages across protocols (Figure S2). To further compare the cellular heterogeneity of primary cortical tissue and organoids, we performed immunohistochemistry for proteins that label radial glia and excitatory neurons. The human and macaque primary cortex samples displayed clear organization of radial glia and intermediate progenitors in the ventricular and subventricular zones and neurons migrating toward the cortical plate over an extensive intermediate zone. Similarly, cortical-like organoids from both human and chimpanzee contained ventricular and subventricular zone-like structures in which cells expressed markers of radial glia and intermediate progenitors (Fig. 1D). Outside of these zones, we observed cells expressing deep and upper layer markers at 6 and 15 weeks of organoid differentiation, respectively (Figures 1D, S1), but the overall distance from the ventricle to the periphery was greatly compressed compared to primary samples.

We next sought to define homologous cell types across model systems and species. Biological and technical differences across primary and organoid cells and between species create challenges for unbiased clustering of combined datasets (Figure S3). To overcome this limitation, we performed canonical correlation analysis (Butler et al., 2018), which finds common sources of variation across datasets, to co-cluster cells from different model systems in three pairwise comparisons. For the organoid report card analysis, we co-clustered human primary and human organoid cells (Figure 2A). For the comparative analysis in primary tissue samples, we coclustered human and macaque primary cells (Figure 2B), and for the comparative analysis in organoid samples, we co-clustered human and chimpanzee cells (Figure 2C). In each case, we identified major telencephalic cell types including radial glia across cell cycle phases, intermediate progenitor cells, excitatory neurons at different stages of maturation, and inhibitory neurons. These cell types were present across model systems and species and were reproducible across individuals and with alternative co-clustering methods (Figures 2A-C, S3), although the relative proportion of these cell types varied between primary sample and organoid models. In addition, we observed that human cells across each of the three comparisons shared similar clustering patterns (Figure S3). Together, these analyses provided a baseline for comparing cells of the same type across model systems and species.

**Figure 2.**
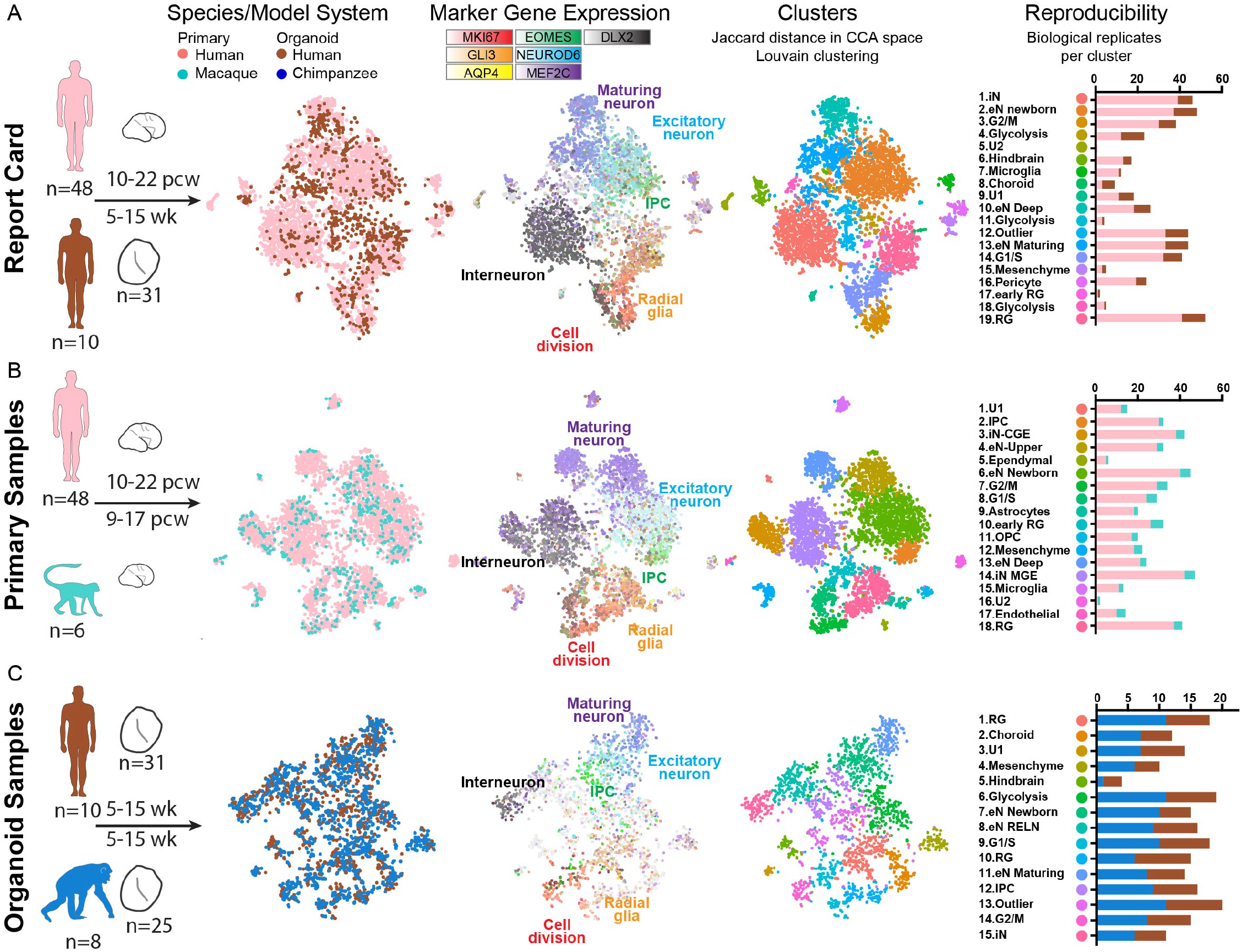
Identification of homologous cell types across species and model system. **a-c**) Pairwise comparisons of human primary and human organoid cells (**a**), human primary and macaque primary cells (**b**), and human organoid and chimpanzee organoid cells (**c**), with the number of distinct individuals and organoids depicted under the schematics. Columns 1-3 display cells plotted based on gene expression similarity after principle components analysis and t-stochastic neighbor embedding, and colored by species or model system (column 1), by marker genes for known cell types (column 2), and by clusters following Louvain-Jaccard clustering (column 3). Column 4 indicates the number of primary individuals or distinct iPS lines contributing to each cluster.

### Preservation and conservation of gene co-expression patterns

Gene co-expression relationships often reflect biological processes related to cell type, cell state, and signaling pathways, in which many genes with related functions are coordinately regulated (Oldham et al., 2008) (Langfelder and Horvath, 2008). To identify gene co-expression relationships, we performed weighted gene correlation network analysis (WGCNA) separately in each dataset (Figure 3A). Importantly, gene coexpression relationships can be determined independently from cell clustering results, providing an additional method for comparing datasets. We applied this approach to examine the extent to which gene co-expression modules are preserved between primary samples and organoid models and conserved across species by separately determining gene co-expression relationships in all primary human cells, all primary macaque cells, all human organoid cells, and all chimpanzee organoid cells (Tables S4-S6). We found that the majority of modules derived independently in each dataset were highly correlated (Pearson’s R > 0.5) (Figures 3A, S4). For example, over 70% of human primary modules showed a Pearson’s correlation greater than 0.7 with a human organoid module. Overall, the shared patterns of co-expression provided systematic evidence for preserved gene regulatory mechanisms in organoid models and an overall conservation of these regulatory relationships across species.

**Figure 3.**
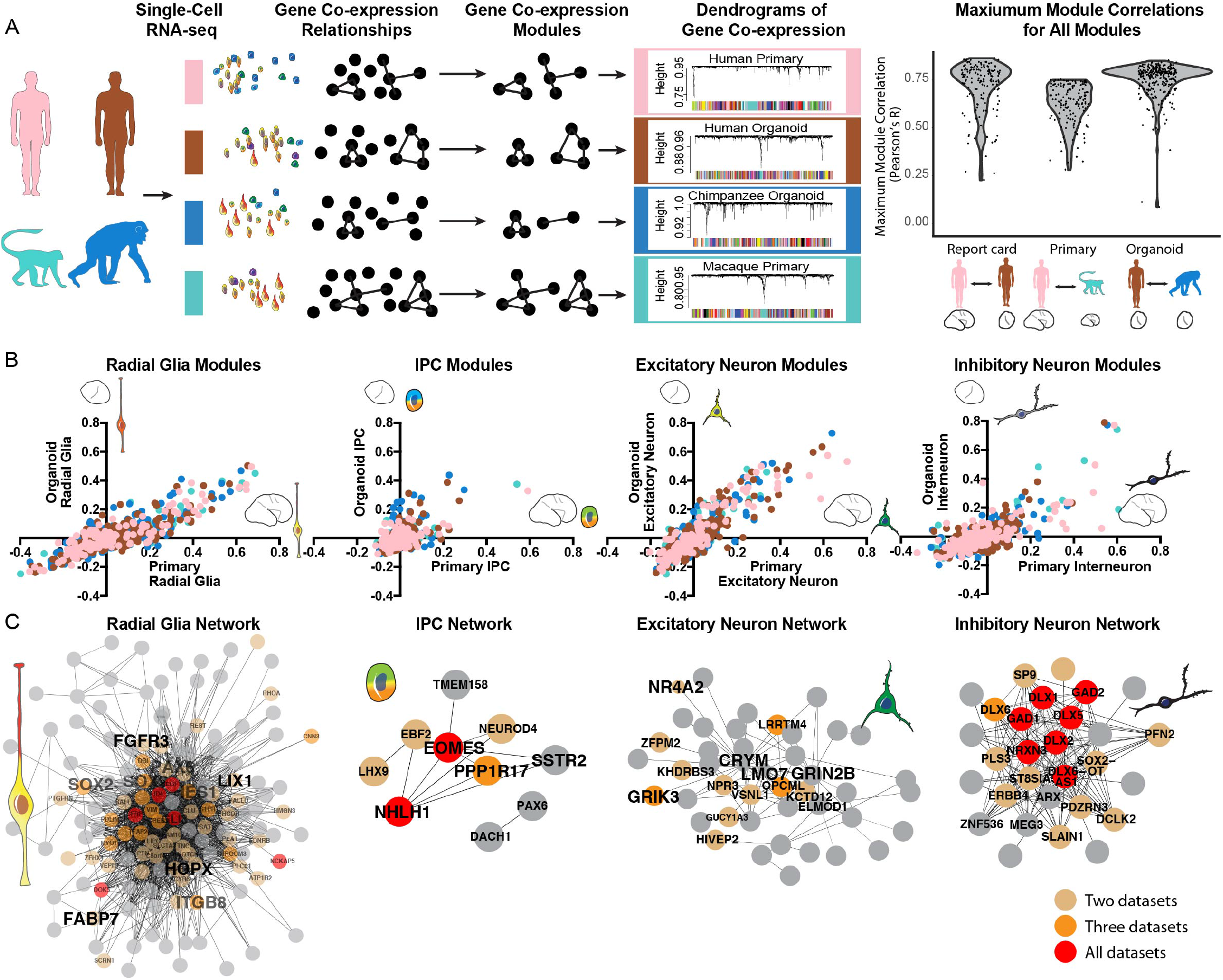
Conservation of gene co-expression modules across species and model system. **a**) Gene co-expression relationships were determined independently in each dataset using WGCNA. Violin plots indicate the distribution of maximum correlation values for all co-expression modules in each pairwise comparison. **b**) Scatterplots depict the correlation of modules to cell types determined in organoid and primary cell datasets. Dots are colored by the model system in which the network was identified. **c**) Network maps depict the correlation of genes from top cell type networks across all four datasets. Edges represent a correlation with R > 0.25, with edge length inversely related to correlation strength. Brown, orange and red dots highlight genes that appear in the core module for 2, 3 and 4 networks respectively.

We next examined whether the highly correlated modules related to common biological processes. Using a general linear model, we identified many gene co-expression modules related to cell type, but relatively few modules related to individual differences, organoid model system, or species differences (Figure S4). For example, many modules generated in each dataset correlated with radial glia and excitatory neuron clusters, while a subset of modules correlated with intermediate progenitor and interneuron subtypes (Figure 3B). These relationships allowed us to generate consensus modules related to cell type that emerge across all datasets (Figure 3C). In addition, highly correlated modules that related to cell states independently emerged in all four datasets, including the G2/M and G1/S transitions and excitatory neuron maturation (Figure S4). On the other hand, modules related to cell types not commonly found in organoids, such as microglia, and oligodendrocyte precursors, commonly emerged across species, but only in primary cell datasets (Figure S4). The independent emergence of modules highly correlated to cell type suggests that transcriptional programs determining cell types are largely conserved among primates and that gene regulatory mechanisms in major telencephalic cell types and states can be modeled in cerebral organoids.

### Developmental processes and metabolic states

We further examined the extent to which organoid models preserve developmental trajectories related to neuronal differentiation, radial glia maturation, and cortical areal identity that we recently described during normal cortical development (Nowakowski et al., 2017). As previously reported, the signature of differentiation from radial glia to excitatory neurons is highly correlated between primary cells and organoid models, with very few genes deviating from the *in vivo* trajectory (Figure 4A) (Camp et al., 2015; Pollen et al., 2014). We similarly found that gene co-expression modules representing radial glia maturation independently arose in primary and organoid radial glia datasets (Figures 4B, S5). In addition, astrocyte production was restricted to the oldest organoid samples (Figure S5) as recently reported (Sloan et al., 2017). However, the radial glia maturation signature was more strongly correlated with primary sample age than with organoid age, indicating variability in the timing of organoid maturation (Figure 4B). The emergence of distinct cortical areas is a major feature of normal cortical development, and maturing excitatory neurons show strong transcriptional signatures of areal identity. Therefore, we used excitatory neurons from human primary visual (V1) and prefrontal cortex (PFC) to construct a classifier for the areal identity of organoid neurons (Figure 4C). Consistent with previous immunohistochemical observations of gradients in several regional identity genes (Eiraku et al., 2008; Kadoshima et al., 2013), most organoids contained a mixture of V1-like and PFC-like excitatory neurons and additional neurons of unknown areal identity (Figure 4C). Overall, neuronal differentiation, radial glia maturation, and aspects of cortical arealization occurred spontaneously in organoid models, but the timing of radial glia maturation and arealization of neuronal identity was heterogeneous.

**Figure 4.**
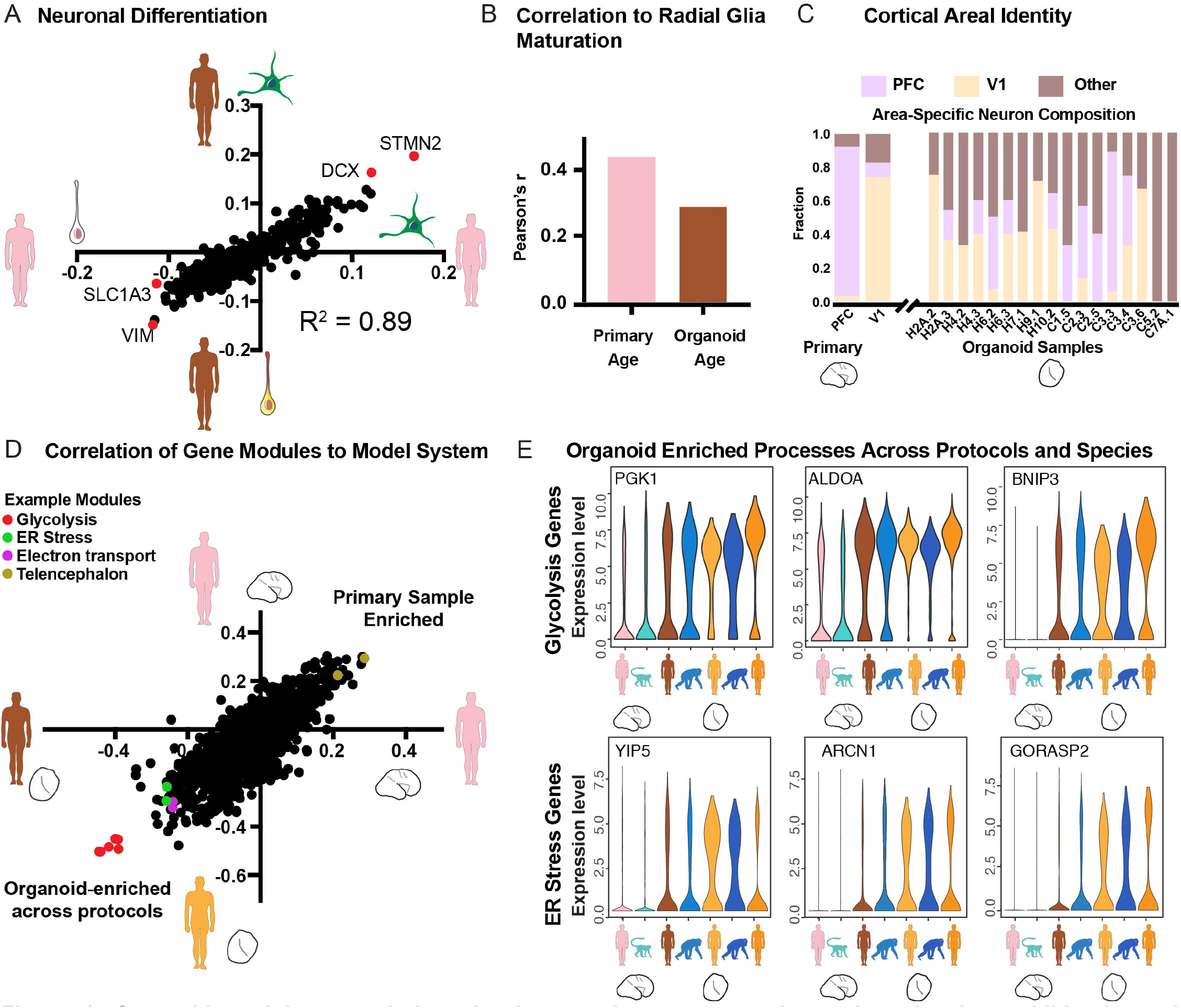
Organoid modules recapitulate developmental gene expression trajectories but exhibit elevated metabolic stress across protocols. **a**) Scatterplot shows the correlation of genes to the neuronal differentiation signature as derived in organoids (Y-axis) and in primary cells (X-axis). **b**) Histogram of R values indicating the correlation of the radial glia maturation network to sample age across primary radial glia (PCW8-22) and organoid radial glia (Week 5-15). **c**) Areal identity of maturing excitatory neurons in primary tissue and across organoids as predicted by a classifier. First two columns indicate primary cells with known areal identity. **d**) Scatterplot shows the correlation of all gene coexpression modules to primary human cells (positive on both axes) versus organoid cells from this paper using the Kadoshima protocol (negative on X-axis) and a previous paper (Camp et al., 2015) using the Lancaster protocol (negative on Y-axis). Modules are generated independently in each dataset and correlated to primary or organoid cell identity. Glycolysis, endoplasmic-reticulum (ER) stress, and electron transport modules show a strong correlation with organoid cells from both protocols. **e**) Violin plots illustrate the distribution of single cell gene expression values for hub genes in the glycolysis and ER stress co-expression modules from primary human and macaque cells (columns 1 and 2) organoid cells generated in this paper using the Kadoshima protocol (columns 3 and 4), and organoid cells generated using the Lancaster protocol (columns 5 and 6) and a spheroid protocol (column 7, Sloan et al., 2017).

We next explored the biological processes that were strongly associated with organoid models when compared to primary cells by considering gene co-expression modules enriched in organoid cells. We hypothesized that differences resulting from the organoid system might be reflected across a range of cerebral organoid protocols and might illuminate general opportunities for improving these culture systems. Indeed, we found that the gene co-expression modules up-regulated in organoids compared to primary cells were shared across three protocols varying in their use of patterning molecules to constrain regionalization (Figure 4D, S5). In particular, the modules with strongest organoid enrichment across protocols related to glycolysis (Figures 4D, S5). Interestingly, similar glycolysis gene co-expression modules emerged in primary human and primary macaque cells, but the expression of genes in these modules was higher and more pervasive in organoid cells (Figure 4E), indicating over-activation of a normal metabolic pathway. In addition, we observed enrichment in organoids for modules involving endoplasmic reticulum stress, including the unfolded protein response pathway and electron transport (Figures 4E, S5). In contrast, modules enriched in primary samples related to telencephalon regional identity and excitatory neuron subtypes that were also commonly observed in organoids but found at a higher frequency in primary samples (Figure 4D). Overall, current organoid models preserve major cell types, gene co-expression networks, and developmental trajectories observed in primary tissue, while up-regulated metabolic stress pathways across major protocols provide specific targets for optimization of future organoid culture systems.

### Human-specific gene expression changes

The preservation of gene regulatory networks and developmental processes in organoid models allowed us to search for gene expression patterns that evolved in the last six million years. Our strategy involved first comparing primary human and primary macaque cortex to identify gene expression differences that occur during normal development across more distant primates. Next, to identify which of these changes occurred in recent evolution, we compared gene expression in human and chimpanzee organoid models. We used a likelihood ratio test (LRT) commonly applied to single cell gene expression data and a general linear model to calculate differential expression, accounting for a large number of technical and biological replicates (Methods, Table S7). Across primary cells, we observed 1258 differentially-expressed genes between human and macaque and across cells from organoids with a telencephalon identity, we observed 738 differentially-expressed genes between human and chimpanzee (Figure 5A, S6). The fold change and direction of these differentially expressed genes was correlated (Figure 5B-C) and 261 genes overlapped across comparisons, providing a set of candidate genes independently supported by primary tissue and organoid comparisons that are likely to have derived human-specific gene expression patterns during cortical development.

**Figure 5.**
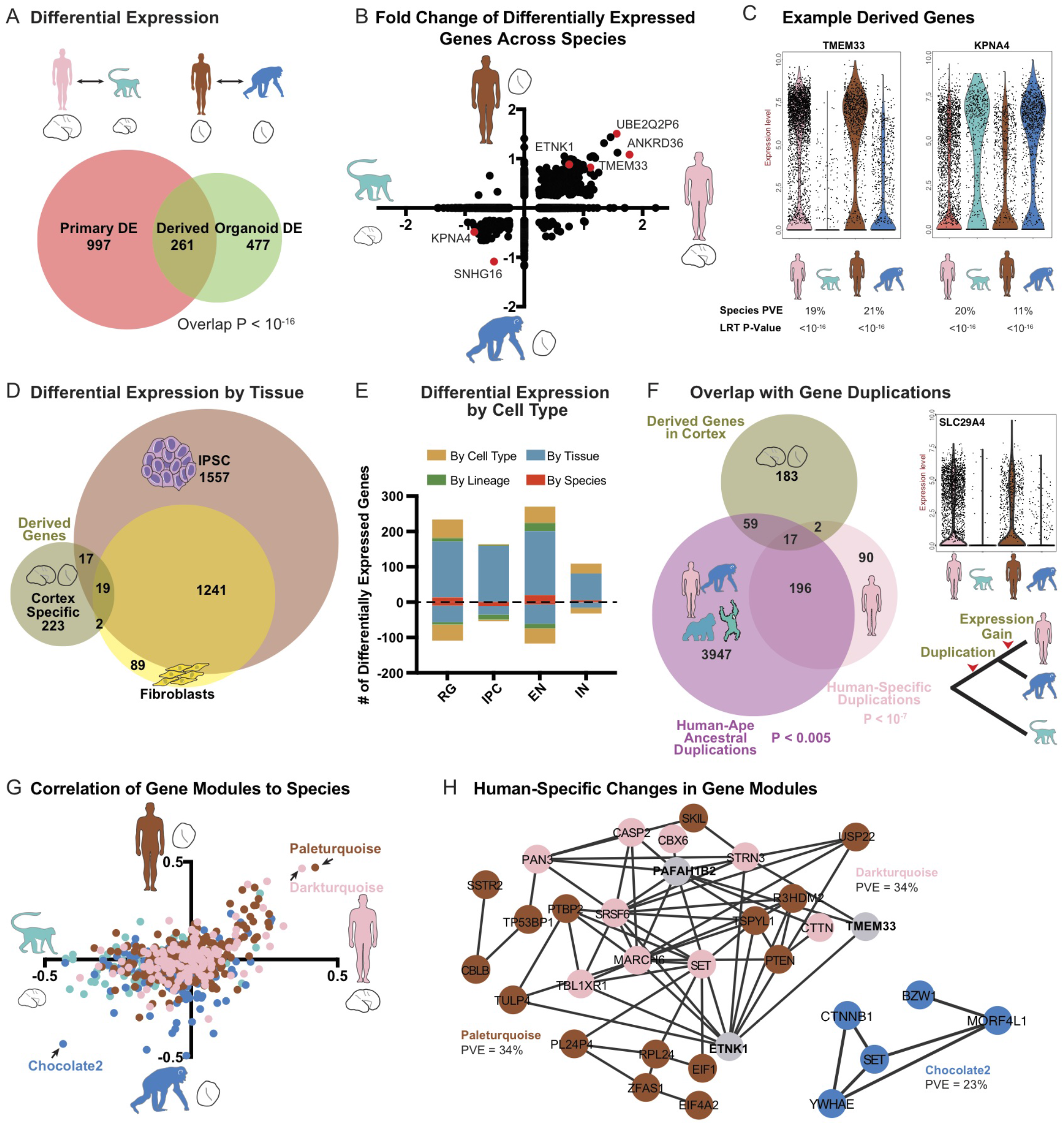
Human-specific gene expression patterns during cortical neurogenesis. **a**) Venn diagram represents the number of differentially expressed genes between primary human and macaque (left circle) and differentially expressed genes between human and chimpanzee organoids with cortical identity (LRT adjusted p-value < 0.0005). Overlap represents candidate genes with human-specific regulatory changes, **b**) Scatterplot illustrates the fold change for genes differentially expressed in either primary cell or organoid cell comparisons across all cells, **c**) Violin plots for genes up- or down-regulated specifically in human cells, **d**) Venn diagram represents the overlap between derived regulatory changes in cortex and human and chimpanzee differential expression previously determined in fibroblasts and pluripotent stem cells (Gallego Romero et al., 2015). **e**) Histogram highlights the number of derived expression changes that are shared across cortex, iPSC, and fibroblasts (by species, red), found only in cortex (by tissue, blue), found only in the excitatory neuron lineage of radial glia, intermediate progenitor cells (IPC) and excitatory neurons (by lineage, green), or in one cortical cell type (by cell type, dark yellow). **f**) Venn diagram represents the overlap between derived genes in cortex, and genes with duplications or copy number expansions that are human-specific (pink) or occurred in apes along the lineage leading to humans, prior to our divergence with chimpanzee (purple) (Sudmant et al., 2013). Note that some genes underwent multiple duplication events along the human lineage (overlap between pink and purple circles). **g**) Scatterplot illustrates the correlation of each gene co-expression module to species across primary cells (X-axis) and organoid cells (Y-axis). Colors correspond to the dataset in which the module was generated. **h**) Network maps highlight genes from the modules with expression most correlated to human or primate cell sources. Edges correspond to a correlation R > 0.25.

We further examined the properties of derived genes with respect to the specificity of differential expression and the overlap with genes underlying neurodevelopmental disorders. To examine the specificity of gene expression changes, we compared our results in developing cortex with recent studies in human and chimpanzee fibroblasts and iPS cells (Gallego Romero et al., 2015). Of the 261 candidate derived genes, 85% of the derived regulatory changes were specific to cortical development (Figure 5D, S6). Although the majority of expression changes were restricted to cortical development, these differences were frequently shared across cortical cell types (Figure 5E, 6S). Because genes linked to autism spectrum disorder also show enriched expression in the stages and cell types of cortical development that we surveyed (Willsey et al., 2013; Parikshak et al., 2013; Bakken et al., 2016), we examined whether derived regulatory changes included disease-linked genes. Of 78 genes with an excess of *de novo* mutations in autism (Stessman et al., 2017), five overlapped with derived expression changes on the human lineage: *PTEN, TRIO, ADNP, TCF7L2*, and *NAA15*, which was modestly significant against a background set of all detected genes (P = 0.004). Thus, our approach of combining primary sample and organoid comparisons revealed many candidate tissue-specific gene expression patterns during human brain development, and highlighted examples of disease-related genes that may have human-specific developmental functions.

Recent studies have identified human-specific genetic changes likely to influence gene regulation or gene copy number (Dennis et al., 2012; McLean et al., 2011; Pollard et al., 2006; Charrier et al., 2012; Dennis et al., 2017). By overlapping major classes of structural genomic changes with human-specific gene expression differences, we sought to identify candidate sequence changes particularly likely to influence gene expression during cortical development. We found that recently duplicated genes (Sudmant et al., 2013) were significantly over-represented among differentially-expressed genes (Figure 5F, P < 10^−7^). For example, nearly all genes in the *SMN1* and *ARL17A* loci showed increased expression in human developing cortex compared to chimpanzee and macaque (Figure S6). This is perhaps unsurprising as the human genome contains additional copies of these duplicated genes. Nonetheless, human-specific duplications may also result in daughter genes with new expression patterns (Dennis et al., 2017). Indeed, we observed examples, such as the NPIP family, that has expanded in copy number in the ape and human lineage (Johnson et al., 2001), in which only a single human paralog, *NPIPB5*, shows derived regulatory changes in cortical tissue (Table S8). Duplications that occurred earlier in ape evolution may also provide a substrate for additional regulatory changes (Dennis et al., 2017; Ohno, 1970). Therefore, we examined whether duplications that occurred in the human lineage prior to our divergence from chimpanzee were also enriched among derived regulatory changes. Overall, we found a modest enrichment for differential expression among these older duplication events (P < 0.005). These include intriguing examples, such as the monoamine transporter *SLC29A4*, which appears to have duplicated in the common ancestor of human and chimpanzee, but only shows expression changes in human cortical development (Figure 5F, S6).

Models of regulatory evolution suggest that genetic changes influencing hub genes may increase the expression of species-specific regulatory networks (Nowick and Stubbs, 2010). To identify gene networks that changed together in the human lineage through the influence of common regulatory mechanisms, we further analyzed whether gene co-expression modules showed human-specific expression patterns. We found that many co-expression modules enriched in primary human compared with primary macaque samples were also up-regulated in human organoid compared with chimpanzee organoid samples, suggesting human-specific regulation of these networks (Figure 5G). The two strongest networks with a derived gain of expression in the human lineage were enriched for negative regulation of transcription related to the G1/S transition and neuronal apoptosis (P < 10^−6^). Together, these results provide candidate regulatory networks that may have evolved together through coordinated transcriptional changes.

Radial glia are particularly likely to influence the evolutionary expansion of the brain (Noctor et al., 2001; Rakic, 2003), and we next examined whether any differentially-expressed pathways might influence radial glial development. Among genes with increased expression in the human lineage, we observed a trend for upregulation of genes in the PI3K/AKT/mTOR pathway (Figures 6A). In particular, insulin receptor (INSR) and PTEN showed strong up-regulation in both human organoids and human primary cells. While this pathway is known to influence neuronal maturation (Takei and Nawa, 2014), we recently described strong immunoreactivity for the mTOR effector phosphorylated ribosomal S6 kinase (pS6) in the outersubventricular zone, specifically in outer radial glia (Nowakowski et al., 2017) indicating that this pathway may also influence outer radial glia during stages of neurogenesis. Given the trend for increased expression of mTOR pathway genes in human cells, we investigated whether the pattern of pS6 phosphorylation during human neurogenesis diverged from that in primates. Using an antibody to conserved phosphorylation sites, we observed comparable levels of immunoreactivity among neurons in the human and macaque cortical plate (Figure S7). However, in germinal zones, we found that pS6 strongly labeled human primary and organoid radial glial cells, in particular those away from the ventricle and ventricular zone-like regions (Figures 6B-C, S6). Conversely, staining in primary macaque and chimpanzee organoid sections showed less pervasive and fainter pS6 immunoreactivity than in human radial glia (Figures 6B-C, S6). Together, these results suggest that changes in the activity of the mTOR signaling pathway have evolved recently in human outer radial glia.

**Figure 6.**
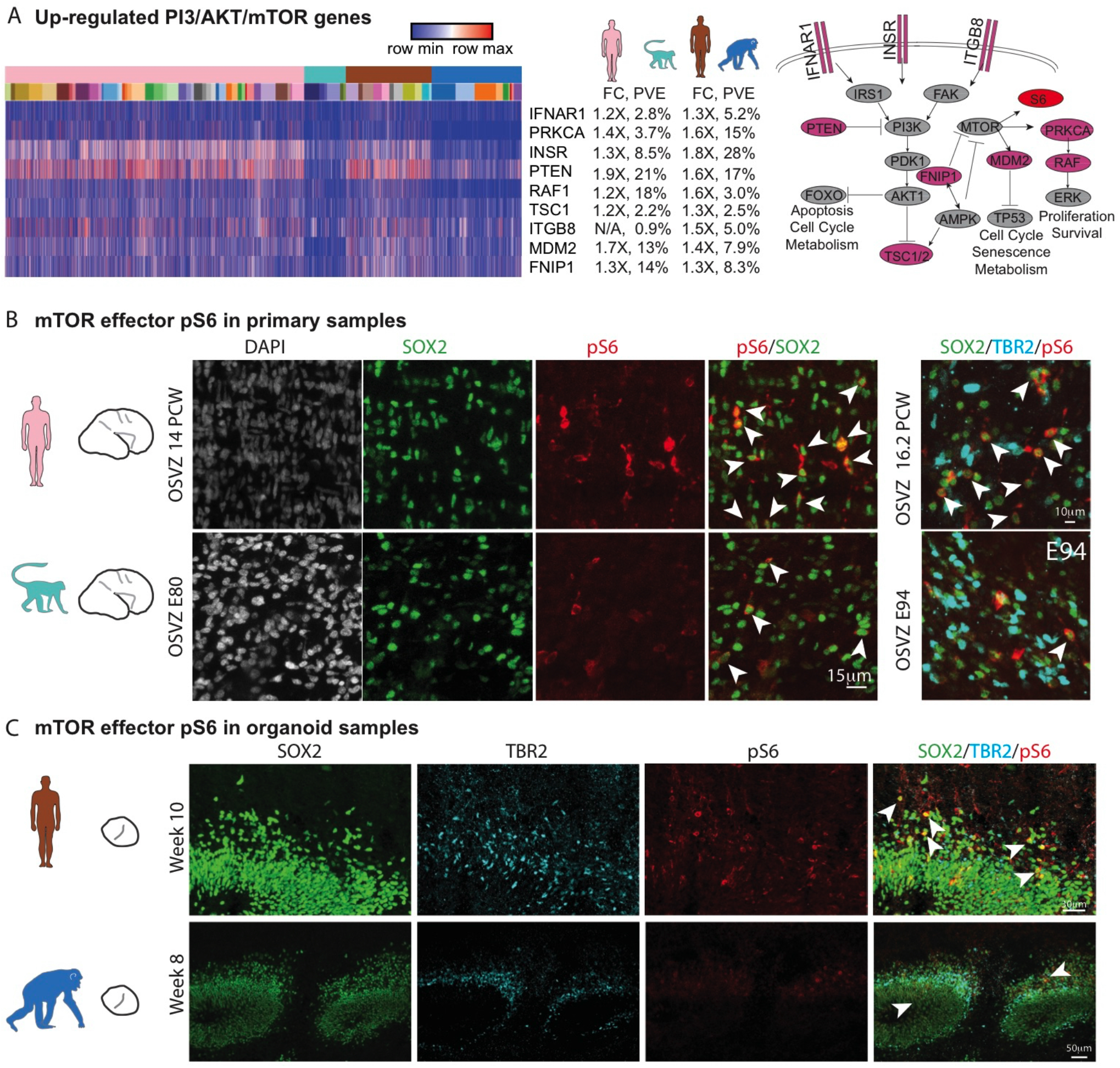
Human outer subventricular zone radial glia show increased phosphorylation of the mTOR effector S6 compared to other primates. **a**) Heatmap illustrating differential expression for genes related to the PI3K/AKT/mTOR pathway, with label showing the log_2_(fold change) and percent variance explained by species in both the primary cell and the organoid cell comparison. Differentially-expressed genes in the context of the PI3K/AKT/mTOR pathway. Genes highlighted in pink are up-regulated in human and phosphorylated S6 (pS6) is marked in red. **b**) Immunohistochemistry illustrates abundant labeling of pS6 in radial glia of the primary human outer subventricular zone compared to the primary macaque cortex. **c**) Immunohistochemistry shows pS6 labeling in radial glia adjacent to cortical-like ventricular zones of human organoid samples which is more abundant than in the chimpanzee organoid. Arrowheads highlight example cells immunoreactive for pS6 and for SOX2. TBR2 (EOMES) labels a separate population of intermediate progenitors.

## Discussion

Over the last six million years, genetic changes along the human lineage have driven remarkable changes in human brain structure, cognition, and behavior. These genetic changes likely influenced patterns of brain development because brain size differences appear *in utero*, and nearly all cortical neurons are generated prior to birth (Sakai et al., 2012). Thus, comparative studies of brain development are essential for understanding the molecular basis of human brain evolution. Genetic changes are particularly likely to influence gene expression, either directly through copy number changes or *cis*-regulatory changes or indirectly through the trans-regulatory effects of genes harboring mutations. However, in contrast to postnatal and adult samples (Liu et al., 2012; Sousa et al., 2017), prenatal brain tissue from chimpanzee, our closest living relative, is largely inaccessible to analyses of gene expression. Here, we sought to establish great ape cerebral organoids as a model system for studying human-specific molecular changes that influence neurogenesis. We found that gene regulatory mechanisms observed during neurogenesis in primary cortex are mostly recapitulated in organoid models, and we identified human-specific gene expression patterns during cortical neurogenesis through independent comparisons of human and primate primary cells and organoid models.

Because cerebral organoids are a relatively new model system, we developed a multi-dimensional report card of the correspondence between human cerebral organoids and primary tissue. Our findings indicate that organoid models across many individuals, species, and protocols reproducibly contain cell types, gene coexpression relationships, and developmental processes homologous to those found in primary cells. The pathways distinguishing organoids - glycolysis, endoplasmic reticulum stress, and electron transport - provide useful targets for further optimization of cerebral organoid media and perfusion conditions and may also have implications for organoid studies of neurodegenerative disorders in which cellular stress pathways are known to accelerate the appearance of disease phenotypes. Together, our report card reveals the fidelity and robustness of developmental programs that can be modeled using cerebral organoids, and serves as a data-driven framework for evaluating organoid cells along biological axes of variation related to cell heterogeneity, neuronal differentiation, radial glia maturation, cortical arealization, and metabolic states. From the report card analysis, we concluded that comparative investigation of organoid models across species provides a realistic window into gene regulatory changes in evolution, but that independent analysis of primary cells from different species still provides crucial validation data independent from organoid models for determining species-specific gene expression patterns.

Human-specific genetic changes may affect cortical development by influencing gene regulatory networks active during neurogenesis. Although there is great interest in modeling the impact of human-specific genetic changes on brain development (Boyd et al., 2015; McLean et al., 2011; Pollard et al., 2006; Prabhakar et al., 2006; Charrier et al., 2012; Dennis et al., 2012; Florio et al., 2016) a description of actual molecular differences during normal development is crucial for interpreting the consequences of genome sequence changes. Against a background of highly conserved gene expression patterns, we identify 261 genes with derived expression changes in the human lineage. A subset of these changes are concerted across non-neural cell types, but the majority appear to be specific to developing cortex and shared across cortical cell types. These results are consistent with a hierarchical model in which regulatory mutations have different levels of pleiotropy from organism to tissue to cell type (Liang et al., 2018). In addition to individual genes, we identified several gene co-expression modules with increased expression along the human lineage, including modules enriched for genes affecting transcription during the G1/S transition and neuronal apoptosis, two developmental processes that may relate to brain expansion by increasing the number of progenitors and neurons. Future analysis of hub genes and candidate upstream factors in these networks may illuminate the causative mutations underlying gene expression differences. In addition, the recent de novo assembly of the chimpanzee genome and improved gene models enabled more accurate measurements of gene expression as a result of higher alignment rates to the native genome of each species (Kronenberg et al., 2018, Fiddes et al., 2018). Future improvements of gene annotations and genome assembly through long read sequencing in other primates including macaque will further strengthen comparative analyses, and increased cellular coverage through droplet capture of single cells across many individuals will make analysis more comprehensive, illuminating the most divergent gene networks and cell types.

Several influential models suggest that human brain expansion may result from an increase in the number of radial glia early in development (Noctor et al., 2001; Rakic, 2003), an increase in the number of outer subventricular zone progenitors (Fish et al., 2008; Kriegstein et al., 2006; Smart et al., 2002), and an extended neurogenic window (Rakic, 1995). In each of these models, signaling pathways controlling radial glia expansion and self-renewal represent candidate molecular pathways that could explain changes in developmental cell behavior (Fietz et al., 2012). We recently observed increased activation of the LIFR/STAT3 self-renewal pathway in outer subventricular radial glia compared to ventricular radial glia, which could support selective expansion of this secondary proliferative population during cortical development. However, this molecular signature was conserved in macaque radial glia and not likely related to more recent human brain evolution (Pollen et al., 2015). Here, our comparative transcriptomic data among primates allowed us to examine additional signaling pathways that may have changed in more recent evolution. We find evidence that an increase in the activity of another key signaling pathway, PI3K/AKT/mTOR may distinguish human outer radial glia from those in other primates. In many stem cell contexts, the mTOR pathway has been shown to promote stemness and long-term self-renewal (Zhou et al., 2009; Rafalski and Brunet, 2011) suggesting that selective upregulation of this pathway in outer radial glia could contribute to human brain expansion by increasing the neurogenic capacity of these cells. Importantly, mTOR pathway mutations are associated with several human disorders, including autism, focal cortical dysplasia, and glioblastoma multiforme (Poduri et al., 2012; Wang et al., 2017; Ceccarelli et al., 2016). Our results suggest that human outer radial glia may be particularly sensitive to perturbations in this pathway, and future studies may more directly assess the role of this pathway in human and primate neurogenesis. Together with recent work, our study adds to a burgeoning field of “cellular anthropology” (Prescott et al., 2015) by providing a comparative organoid platform for more systematically characterizing the specialized molecular features of human cortical development and evolution.

## Supplementary Tables

**Table S1. iPS Lines** Details of iPS cell lines used in this study, including cell line origin, clone name, reprogramming method, and protocol used for differentiation.

**Table S2. Cell Metadata** Relevant metadata attributes of all cells analyzed in this study.

**Table S3. Cluster Assignments and Interpretations** For each cell, the cluster identity in each homologous cell type analysis is noted and the interpretation of each cluster based upon key marker genes is presented.

**Table S4. WGCNA Gene Modules** The modules independently generated are shown by gene, with the gene, the module membership and the dataset of origin described.

**Table S5. Module Eigengene Values** Module eigengene values for all networks used in this study and their module eigenvalues for all cells analyzed.

**Table S6. Module Correlation to Property** The correlation value and percent variance explained of each gene coexpression module to a property of interest including cell type, species, and protocol is presented.

**Table S7. Percent Variance Explained (PVE) By Genes** For all the genes used in the analysis, the percent variance explained by Cell Type, Donor ID, and Species is shown for both primary and organoid species comparisons.

**Table S8. Derived Genes** Intersections of the individual analyses exploring differential gene expression between species is shown with derived up genes indicating genes that are up-regulated in human and derived down genes indicating genes that are down-regulated in human. NHP refers to non-human primate, in this case referring to the intersection of genes upregulated in both macaque and chimpanzee compared to human.

**Table S9. Differential Gene Expression** The full set of differential expression analysis in both primary human versus primary macaque samples and human versus chimpanzee organoids. The comparison is performed in a concerted fashion as well as separately in radial glia, intermediate progenitor cells, excitatory neurons and inhibitory neurons.

## ACKNOWLEDGMENTS

The authors thank Y. Gilad, T. MacKenzie, S. Mokhtari, A. Tarantal, A. Fox, S. Wang, L. Adame, C. Villarreal, G. Wang, M. Snyder, T. Wang, D. Gordon, D. Kingsley and members of A.R.K and A.A.P labs for helpful discussion, reagents, and samples. This study was supported by CIRM (GC1R-06673-C to A.R.K.), Damon Runyon Foundation (DRG-2166-13 to A.A.P.), NIH (U01 MH105989 to A.R.K, F32 NS103266-02 to A.B.), Simons Foundation (SFARI 491371 to TJN) the Bowes Foundation, and ORIP/OD P51OD011132.

## Author contributions

Methodology: AB, TJN JS, AAL, JAW, CL, OSM, AP. Investigation: AAP, AB, TJN, OSM, EDL, MAMR, BA, MaB, MeB, MGA. Resources: BP, MaB. Software: CBL, ITF. Formal analysis AB, TJN, ZNK, MLD, CBL, ITF, OSM, AAP. Writing: AAP AB, with input from all authors. Funding acquisition ARK AAP. Conceptualization AAP. Supervision: ARK AAP SRS, DH, EEE.

## Methods

### Cell lines and Samples

We used previously described human and chimpanzee iPSC lines (Bershteyn et al., 2017; Gallego Romero et al., 2015; Pavlovic et al., 2018), and generated four additional lines from three chimpanzees (Table S1). All new lines were reprogrammed from fibroblasts using episomal plasmids according to a recently published protocol (Okita et al., 2013) and matching the protocol used for the majority of existing human and chimpanzee lines used in this study (Table S1). Low passage fibroblasts (P3 – P7) were obtained from the Coriell Cell Repository (Pt1: 12 year old chimpanzee male, catalog: PR00226; Pt2: 6 year old chimpanzee male, Maverick, catalog: S003611; Pt5: 8 year old chimpanzee male, catalog PR00738). For each line, we electroporated 300,000 fibroblasts with three micrograms of an episomal expression plasmid mixture encoding OCT3/4, SOX2, KLF4, L-MYC, LIN28, and shRNA for TP53 using a Neon Electroporator (Invitrogen), and a 100 μL kit, with setting of 1,650V, 10ms, and three pulses, as previously described (Bershteyn et al., 2017). After 5 – 8 days, electroporated cells were detached and seeded onto irradiated SNL feeder cells. Culture medium was replaced the next day with primate ESC medium (Reprocell) containing 5 – 20 ng/mL of βFGF, with higher levels of FGF producing better results. After 20 – 30 days, colonies were picked and selected for further cultivation. After three to five passages, colonies were transferred to Matrigel-coated dishes and maintained in mTeSR1 medium (Stem Cell Technologies, 05850) supplemented with Penicillin/Streptomycin/Gentomycin. Further passaging was performed using calcium and magnesium free PBS to gently disrupt colonies. Each line showed a normal karyotype between passage 10 and 15. Macaque cortical tissue was generously provided from samples being used for other experiments from the UC Davis Primate Center. De-identified tissue samples were collected with previous patient consent in strict observance of the legal and institutional ethical regulations. Protocols, cell lines and samples were approved by UCSF GESCR (Gamete, Embryo, and Stem Cell Research) Committee.

### Organoid and Primary Samples

We generated cerebral organoids according to previously published protocols (Kadoshima et al., 2013; Bershteyn et al., 2017). Human and chimpanzee iPSCs were dissociated to single cells with Accutase and reaggregated in lipidure-coated 96-well V-bottom plates at a density of 10,000 cells per aggregate, in 100 μL of cortical differentiation medium per well. The cortical differentiation medium (Glasgow-MEM, 20% KSR, 0.1mM NEAA, 1mM sodium pyruvate, 0.1mM β-ME, 100 U/mL penicillin/streptomycin) was supplemented with Rho Kinase Inhibitor (Y-27632, 20 μM, Tocris, Cat# 1254 day 0 and day 3), WNT inhibitor (IWR1-ε, 3 μM, Cayman Chemical Cat# 13659 days 0-18) and TGF-β inhibitor (SB431542, Tocris, Cat #1614, 5 μM, days 0-18). Media was changed on days 3 and 6 and then every 2-3 days until day 18. Aggregates were then transferred to ultra low adhesion 6-well plates in DMEM/F12 medium with Glutamax supplemented with N2, Lipid Concentrate, Fungizone (2.5 μg/mL), and penicillin/streptomycin (100 U/mL) and grown under 40% O2 5% CO2 conditions. After five weeks, FBS (10% v/v), Matrigel (1% v/v) and heparin (5 μg/mL) were added to the medium, and after 8 weeks organoids were transferred to lumox dishes (Sarstedt), which have a gas permeable base.

### Immunohistochemistry

Organoids were fixed using 4% paraformaldehyde (PFA) in PBS for 20 min. Primary samples were fixed in 4% PFA prepared in calcium and magnesium free phosphate buffered saline (PBS) (pH~10) overnight at 4°C with constant agitation. After fixation, organoids and primary samples were washed in PBS (pH7.4), equilibrated in 30% sucrose in PBS overnight at 4°C, embedded in blocks with a 1:1 mixture of 30% sucrose/OCT compound (Tissue-Tek, VWR) and frozen at −80°C. Blocks were sectioned to a thickness of 16-20 μm cryosections, and antigen retrieval was performed, by heating sections to 95° in 10mM sodium citrate (pH=6.0) for 15 min. Sections were then permeabilized and blocked with 10% donkey serum in PBS in 2% Triton X-100, 0.2% gelatin. Primary antibody incubations were performed at 4°C overnight and secondary incubations were performed at room temperature for 1-3 hr, followed by three 20 min washes in PBS, and staining with AlexaFluor secondary antibodies (Invitrogen, 1:1000 dilution). Primary antibodies included: chicken anti-TBR2 (EOMES, 1:150, R&D AF6166), rabbit anti-pS6 (1:100, Cell Signaling 2211S), goat anti-SOX2 (1:200, Santa Cruz SC17320), mouse anti-SOX2 (1:200, Santa Cruz SC365964), Mouse anti-TUJ1 (β III TUBULIN, Covance, MMS-435), rabbit anti-PAX6 (Covance, PRB-278P), rat anti-CTIP2 (BCL11B, 1:500, Abcam ab18465), rabbit anti-TBR1 (1:500, Abcam ab31940), rabbit anti-SATB2 (1:1000, Abcam ab34735).

### Single cell RNA sequencing

For single cell dissociation, primary macaque and organoid samples were cut into small pieces, and incubated with a pre-warmed solution of Papain (Worthington Biochemical Corporation) prepared according to manufacturer’s instructions for 10 min at 37°C. After approximately 30 – 60 min incubation, samples were triturated and macaque samples older than E100 and organoids older than 14 weeks were spun through an ovomucoid gradient to remove debris. Cells were then pelleted at 300g and resuspended in PBS supplemented with 3% fetal bovine serum (Sigma). Samples were diluted to approximately 170,000 cells per ml before processing capture using Fluidigm C1 auto-prep system as previously described (Nowakowski et al., 2017; Pollen et al., 2014; 2015). We performed library preparation using the Illumina Nextera XT library preparation kit, and quantification using the Bioanalyzer (Agilent) according to manufacturer’s protocols. Paired-end 100 bp sequencing was performed on the Illumina HiSeq2500.

### Alignments and gene models

Trim Galore 3.7 was used to trim 20 bp of adaptor sequence, and paired-end alignments were performed using HISAT2 to the human reference genome GRCh38, the updated chimpanzee reference genome panTro6 (Kronenberg et al., 2018) or the macaque reference genome, rheMac8. For each cell, counts were determined using the subread-1.5.0 function featureCounts, and counts were normalized to counts per million. To determine orthologous genes across species, whole genome alignments between GRCh38 and either the current primate references (panTro4, gorGor4, ponAbe2, rheMac8) or the updated great ape references panTro6 (Kronenberg et al, 2018) were generated using progressiveCactus (Paten et al., 2011). Outgroup genomes were gibbon (nomLeu3), bushbaby (otoGar3), squirrel monkey (saiBol1) and mouse (mm10). These alignments were then used as input to CAT, using the GENCODE V27 annotation of GRCh38 as the annotation input. Species-specific RNA-seq and IsoSeq were used to help guide the annotation process (Fiddes et al, 2018). For this study, the resulting annotations on rheMac8 and the updated chimpanzee (panTro6) were used. CAT automatically defines orthology relationships using the information inherent in the whole genome alignment as well as subsequent filtering. This allows for quantification to be performed on the native species genome for a given experiment, and the resulting gene counts can be compared across species by their unique gene identifiers. In total, 49,360 genes annotated in human were identified in both chimpanzee and rhesus.

### De-multiplexing chimeric organoids

To identify the species of origin for each cell from RNA-seq data, we compiled a list of diagnostic loci where the chimpanzee and human genome assemblies differ by a single-base substitution mutation that shows no sign of being polymorphic. We then analyze the RNA-seq reads from each cell at these genomic loci and calculate the number of times that we observe the human or chimp base. We created the diagnostic SNPs by examining whole-genome alignments of human (hg19) against chimpanzee (panTro4), gorilla (gorGor3), orangutan (ponAbe2), and rhesus (rheMac3) (Casper et al., 2018). To conservatively filter for syntenic alignments, we removed alignments involving segments that have not been placed on chromosomes or that represent alternative haplotypes as well as mandating similarity scores of at least 10000 (Kent et al., 2003). With the remaining set of high quality alignments, we located positions in the human genome where chimpanzee had a different base from human and there was no current data to suggest the position was polymorphic in dbSnp (Sherry et al., 2001). To guard against highly variable sites or alignment ambiguity, we ensured that at least one outgroup species (gorilla, orangutan, and rhesus) was present and the outgroup species agreed with either chimpanzee or human. This resulted in approximately 15 million diagnostic positions. For each singlecell RNA-seq library we mapped the reads to the human genome and analyzed the reads that overlapped a diagnostic position. In total, three organoids were initially seeded as chimeric in a pilot common garden experiment, but resulted in nearly pure outcomes for one species: H1.C7B.2 had 30/33 human cells, C1.1 had 39/49 chimpanzee cells, and C1.7 had 47/47 chimpanzee cells. As such, these organoids are displayed in Figure 1 and S1 heatmaps by the dominant species. The software used in this analysis is freely available under a BSD-style license (http://www.github.com/craiglowe).

### Clustering and Determining homologous cell types

Datasets underwent quality control individually, removing any cells with fewer than 1000 genes/cell, greater than 20% of reads towards mitochondrial genes, and greater than 10% of ribosomal genes. Each dataset was then normalized and scaled. Clustering of individual datasets was performed as previously described (Lefebvre, 2008; Shekhar et al., 2016; Nowakowski et al., 2017) using a Jaccard weighted nearest neighbor distance in the space of significant principle components followed by Louvain clustering. Homologous cell types were generated using canonical correlation analysis (CCA) (Butler et al., 2018). CCA was performed in the space of 20 correlated dimensions, and tSNE was run in the space of the CCA dimensionality reduction for visualization. Clustering of combined datasets was then performed using the Louvain-Jaccard method in the space of CCA dimensions. We also developed an alternative approach to co-cluster cells across model system or species that involved minimal transformation using a restricted set of marker genes for analysis. We first performed Louvain clustering using Jaccard distance on each dataset separately, as described above. We next selected the top 40 positive markers for each cluster from each dataset based on average difference. We then used a general linear model to remove markers with a high variance explained (typically greater than 0.1% to 1%) by model system or species, modeling these parameters as fixed effects. We then performed clustering of the combined dataset, scaled and normalized together, in the space of these restricted marker genes. This approach did not fully remove batch effects, but also retained cell subtype nuances from the original clustering and provided an independent validation of cell types as determined by CCA. Visualizations were performed in either tSNE space or using violin plots that reflected the log_2_(counts per million) value of gene expression.

### Determining co-expression modules

WGCNA was performed using the top 2000-5000 PCA loading genes ranked across significant PCs based on absolute value of rotation scores using the WGCNA R package (Langfelder and Horvath, 2008). Parameters such as softPower, deepSplit, and cuttreeDynamic were determined independently for each dataset. To identify homologous modules from independent datasets, we used gene coexpression module kME scores across all genes to correlate modules derived from different datasets. Gene network maps presented are based on Pearson’s correlation of expression levels of genes from each module, as specified in each figure legend.

### Differential expression and enrichment analysis

Correlation analysis of networks to cell, species or protocol identity was performed by binarization of the variable of interest and using a Pearson’s correlation to quantify the similarity across the dataset. When a correlation was performed across two cell types or other defined variables across the axis, only this set of cells was used, otherwise the whole dataset was utilized. Differential expression analysis at the gene level was performed using the likelihood ratio test and with a binary distribution suited to zero inflated data. In order to be considered as a differentially expressed gene, the average fold change was required to exceed an absolute value of log2(0.25) and the frequency of expression was required to be greater than 25% across cells in at least one dataset. For each pairwise comparison, we used a general linear mixed model through the variance partition package (Hoffman and Schadt, 2016) to evaluate the extent to which fixed effects of cell type, donor, and species or protocol explain variation in each co-expression network. In all three pairwise comparisons, the majority of networks that could be explained by these factors related to cell type, with donorID, protocol, and species, explaining substantial variation in a smaller set of co-expression modules. Enrichment analysis for module signaling pathways was performed by intersecting gene lists with curated pathways in Enrichr (Chen et al., 2013). Adjusted p-values from this analysis are represented by −log_10_(p-value) in histograms.

